# Long SARS-CoV-2 nucleocapsid sequences in blood monocytes collected soon after hospital admission

**DOI:** 10.1101/2020.12.16.423113

**Authors:** Nathan Pagano, Maudry Laurent-Rolle, Jack Chun-Chieh Hsu, the Yale IMPACT Research Team, Chantal BF Vogels, Nathan D Grubaugh, Laura Manuelidis

## Abstract

Many viruses infect circulating mononuclear cells thereby facilitating infection of diverse organs. Blood monocytes (PBMC) are being intensively studied as immunologic and pathologic responders to the new SARS-CoV-2 virus (CoV19) but direct evidence showing CoV19 in monocytes is lacking. Circulating myeloid cells that take up residence in various organs can harbor viral genomes for many years in lymphatic tissues and brain, and act as a source for re-infection and/or post-viral organ pathology. Because nucleocapsid (NC) proteins protect the viral genome we tested PBMC from acutely ill patients for the diagnostic 72bp NC RNA plus adjacent longer (301bp) transcripts. In 2/11 patient PBMC, but no uninfected controls, long NCs were positive as early as 2-6 days after hospital admission as validated by sequencing. Pathogenic viral fragments, or the infectious virus, are probably disseminated by rare myeloid migratory cells that incorporate CoV19 by several pathways. Predictably, these cells carried CoV19 to heart and brain educing the late post-viral pathologies now evident.

## INTRODUCTION

Some acute lytic viral infections release free viral particles into the bloodstream where they are easily assayed with modern molecular techniques. However, a large variety of viruses travel within white blood cells such as retroviruses, e.g., HIV (*1, 2*), and flaviviruses, e.g., Dengue (*3*). Some viruses carried in white blood cells are associated with chronic disease. Poliovirus, a “neurotropic” enterovirus originally thought to spread directly through nerves, instead transits from gut to white blood cells to spleen, and only later infects brain (*4*). There are at least 13 different classes of DNA and RNA viruses that infect peripheral blood mononuclear cells (PBMC) that can be a reservoir for persistence (*5*) in addition to the infectious Creutzfeldt-Jakob Disease (CJD) agent (*6*). Like HTLV1, a retrovirus that can be sequestered in brain myeloid microglia, the CJD agent in white blood cells progressively increases after primary infection of gut dendritic (myeloid lineage) cells to later show up in highly infectious myeloid microglia of brain (*7, 8*). Experimentally, the CJD agent requires cell-to-cell contact for infection (*9*), probably via viral synapses, the same mechanism or conduit used for T to myeloid cell transmission of HIV (*1*). Moreover, chronic HIV infection of brain microglia is linked to neurocognitive disorders and dementia, as is the very different ∼20-25nm CJD infectious particle (*10, 11*). These observations emphasize the general principle that infected migrating myeloid cells can take up residence in and perpetuate infection and chronic pathology in the brain and other organs. The presence of nucleocapsid CoV19 RNA, especially in migrating myeloid cells, might explain some of the ensuing brain and heart pathologies that were predictable and that have become increasingly evident during this pandemic Coronavirus 2019 Disease (COVID-19).

We were aware that PBMC could contain extremely low CoV19 RNA from infection limited to only rare myeloid cell types. Those cell types can be less abundant than the 1-2% dendritic cells in a PBMC population that is typically dominated by lymphocytes (70-90%). Many different myeloid cell-types are increasingly appreciated and classified, and many of them show wide phenotypic flexibility (*12*). Moreover, total cellular RNA is very low in PBMC, ∼1/40th of that produced by epithelial and cultured tumor cells such as HeLa cells and NIH/3T3 cells. Nevertheless, there were three additional reasons to pursue this study. First, coronaviruses are complex, and can elicit immune responses that damage brain. For example, the mouse hepatitis coronavirus (MHV) induces a post-viral autoimmune demyelinating disease that is a model for Multiple Sclerosis, and other coronavirus strains, such as CoV19, might elicit a different type of post-viral brain pathology. Indeed, prior to the evolution of CoV19, related respiratory coronaviruses that cause human colds have been neuroinvasive, with viral RNA demonstrated in brain parenchyma as well as in myeloid microglia in culture (*13*).

We proposed to study CoV19 in PBMC in April 2020 because little was known about potential immediate viral or late immunologic CoV19 effects on the brain. We suspected that a subset of blood monocytes, not just neural olfactory spread, might be a conduit for CoV19 into brain with subsequent development of neuropsychiatric symptoms. A wide variety of neurologic complications and neuropathology have now been published, e.g., thromboemboli, infarction, radiologic changes consistent with an autoimmune encephalitis (*14)* and even the presence of CoV19 in neurons (*15*). Additionally, routine autopsies of lethal sudden deaths during uncharacterized winter “viral” infections can show classical viral acute lymphocytic and myeloid infiltrates in a normal heart and the recent collapse of healthy young athletes appears to be associated with CoV19 vascular dissemination with microthrombi in the heart and/or brain. Finally, LM had the opportunity to join a zoom meeting at Yale with doctors from Wuhan who shared their data and experience with us early in 2020. One slide showed a striking interstitial pneumonia dominated by many cells consistent with a myeloid lineage. This further encouraged us to find if PBMC infection could be the source of multiorgan spread. We here identify >300nt of CoV19 nucleocapsid RNA in PBMC. These cells can be a conduit for dissemination of the virus, or viral elements, to other organs. Myeloid cells can use several known pathways for CoV19 uptake facilitating its protection and pathological persistence.

## RESULTS

PBMC from 7 male and 7 female patients ranging from 24-82 years old and 4 random uninfected control volunteers were evaluated. The 14 patient PBMC were collected 2-6 days after hospital admission with positive CoV19 swab test. To avoid RNA extraction losses in low RNA PBMC, exhaustive DNA removal was not pursued. PBMC recovery can vary, especially in patients with other underlying illnesses. Thus, all PBMC RNA extracts (from control and CoV19 positive patients) were tested for GAPDH by RT/qPCR to assess amounts of RNA in each; 11 patient samples were adequate for further nucleocapsid (NC) assays, and showed a mean of 69% (±35%SD) of the “normal” controls. The other 3 patients yielded only 1/10^th^ the control RNAs and were insufficient for analyses. CoV19 nucleocapsid RNA was initially evaluated with the standard clinical CDC:N1 primer pair/probe that amplifies a 72bp sequence beginning 14nt downstream from the NC start codon (in red, Table I, Methods). None of the normal samples or water controls were positive. In contrast, three positive RNA controls from 1) the complete RNA nucleocapsid genome (designated G), 2) the complete viral RNA sequence (input at e3 copies) and 3) CoV19 infected Vero cell RNA (designated V+) all produced their expected threshold cycles (Ct) and dilution curves. One patient’s PBMC (18*) was unambiguously positive with Cts of 35.37, 34.02 and 34.91, equivalent to 114, 264 and 152 viral copies respectively (mean of 177) per 5e4 PBMC. This indicated this 72nt NC RNA was present in a low percentage of PBMC. One other patient sample tested in parallel (14*) was ambiguous in this test with a Ct of 37.19 in one of three replicates. All other patient samples were negative for the 72nt RNA. We repeated these 72bp RT/qPCRs in duplicate, and only patient 18* was comparably positive with Cts of 34.2 and 34.7.

**Table I:**
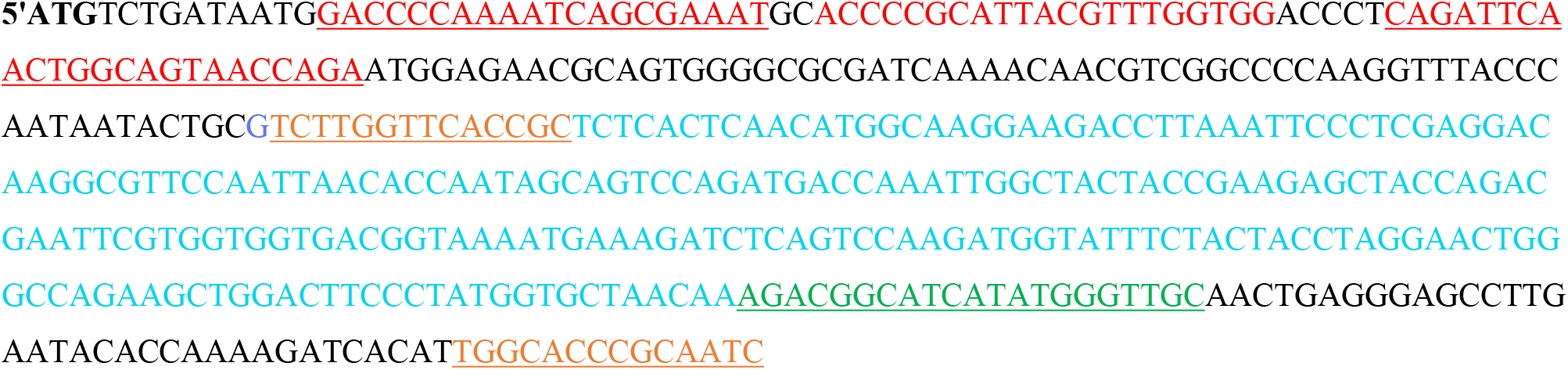
NC Amplification Sequences NC amplified sequences in color downstream from the MET start codon and shows the standard 72 primers and sequence specific FAM probe in red. The 251 and 301bp products sequences are colored with respective primer pairs underlined in different colors. R3 is in green and R5 in orange. The 251bp product is in aqua and the 301bp longer product ends at R5. These amplified RT/qPCR products encompass a substantial (33%) portion of the entire NC RNA that codes for the 455aa nucleocapsid protein.

The 4 control normal samples (designated 1n-4n), and the 11 patient samples (all designated by an *), were subsequently tested for longer adjacent stretches of the NC that covered an additional 301bp as detailed in Table I (see Methods). A single forward primer (F3) was used to generate both a shorter 251bp sequence (with primer R5) as well as a 301bp sequence (with primer R3). This allowed sequencing from two different PCR reactions for validation.

Melting profiles of RT/qPCR products with these primer pairs, using serial dilutions of positive controls along with H_2_O negative controls, revealed significantly different melting temperatures (Tm) for each primer pair product. Fig. 1 shows the clearly different Tm profiles of the 72bp versus the 251 and 301bp amplifications. The CDC:N1 primers melted at 81.45^°^C (±0.07 SD) and had a broad profile. In contrast, the 251bp F3/R5 product gave a significantly higher and sharper Tm of 83.7^°^C (± 0.06 SD). As expected, the longest 301bp product had the highest Tm (84.19^°^C±0.13 SD) with a very sharp and unambiguous curve. This allowed clear identification of real positives from background primer-dimers and other nonspecific artefacts by their Tms. In the CDC:N1 and F3/R3 graphs of standards, the water negative control shows no true melt and only a single profile was obtained. However, in less purified samples Tm peak artefacts were sometimes seen and these typically occurred in PBMC RNA extracts that contained DNA. These nonspecific and variable peaks sometimes precluded simple valid digital Cts. We therefore used the following criteria to validate CoV19-specific peaks: 1) a Tm or peak profile of the PBMC RT/qPCR that matched the positive controls, 2) a minimum of both a positive Tm peak and positive gel band of correct length verified in ≥2 separate aliquots of each patient RNA sample, and 3) sequencing of patient gel bands that matched those from bands of parallel controls including the nucleocapsid genome (G), the e3 reference virus, as well as CoV19 infected Vero cell RNA (V+). Uninfected Vero cell RNA (V-) was also used as a negative total cell RNA control.

**FIG. 1:**
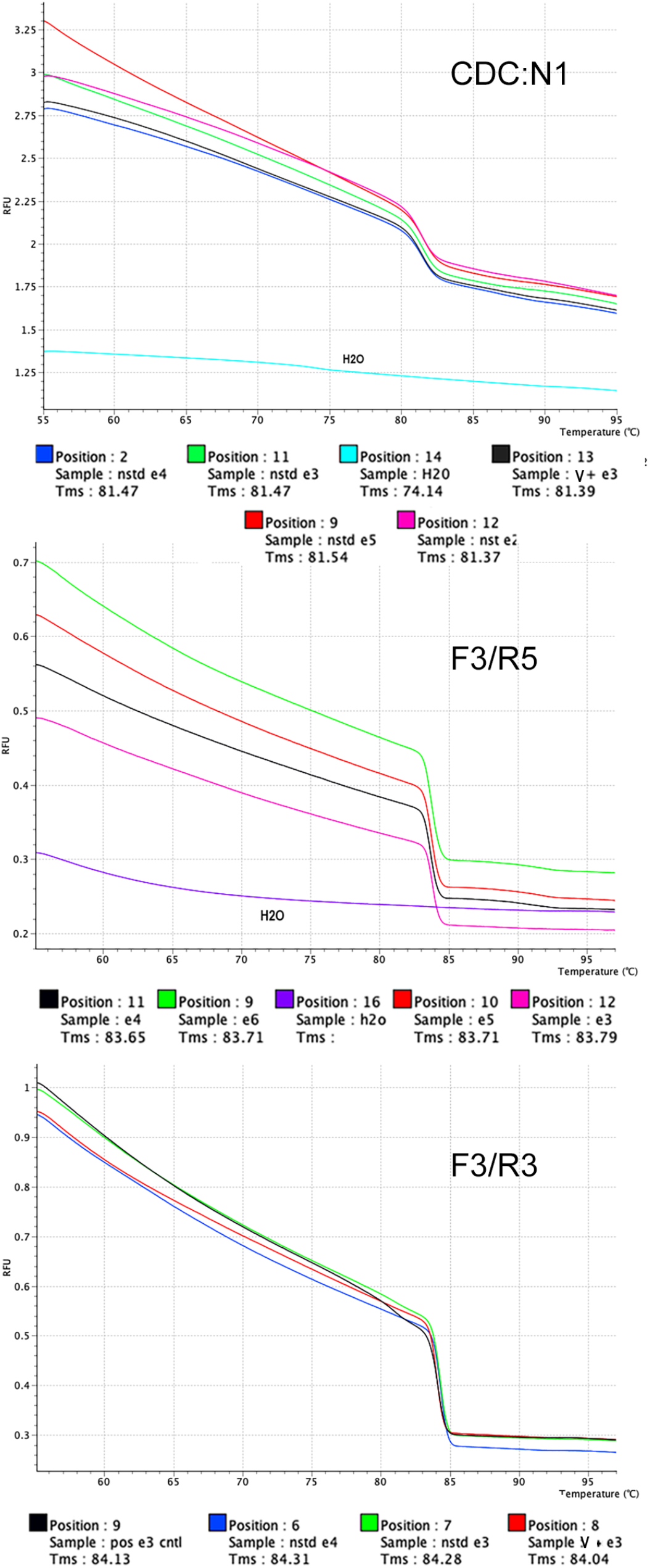
Melt temperature (Tm) of RT/qPCR nucleocapsid (NC) products using different NC primer pairs. The short CDC:N1 product of 72bp has the lowest Tm while the longest F3/R3 pair (301bp) has the highest Tm. Each primer pair Tm product is significantly different (see text).

Fig. 2 shows representative gel runs of RT/qPCR bands for the 251bp (**A**) and 301bp (**B**) NC products. In Fig. 2A only the + lanes show the predicted band, i.e., the nucleocapsid genome control (lane G), the infected Vero cell RNA (V+), and patient extracts 14* and 18*. The second 18* gel band shown is also positive, and from a different aliquot that was run with it’s parallel H_2_O (w) control (lanes 13-14). The 14* band at 251bp is weak but visible, and shows a smear of background that is consistent with non-DNA. Higher loads of this sample on another gel showed a very strong 251bp band (data not shown). Moreover, the RT/qPCR peak profile of the 14* amplified product shown above also revealed an overwhelmingly dominant peak with a Tm matching the other positive controls (see Fig. 3A). Faint smears near the top of the gel in other PBMC samples (lanes 7-10) also indicate large genomic DNA products. The control normal PBMC (n1, n4), uninfected Vero cells (V-), water (w) and other patient samples (11*, 12*, 22*, 23*) are all negative without nonspecific smears. However, some do show primer-dimers.

**FIG. 2:**
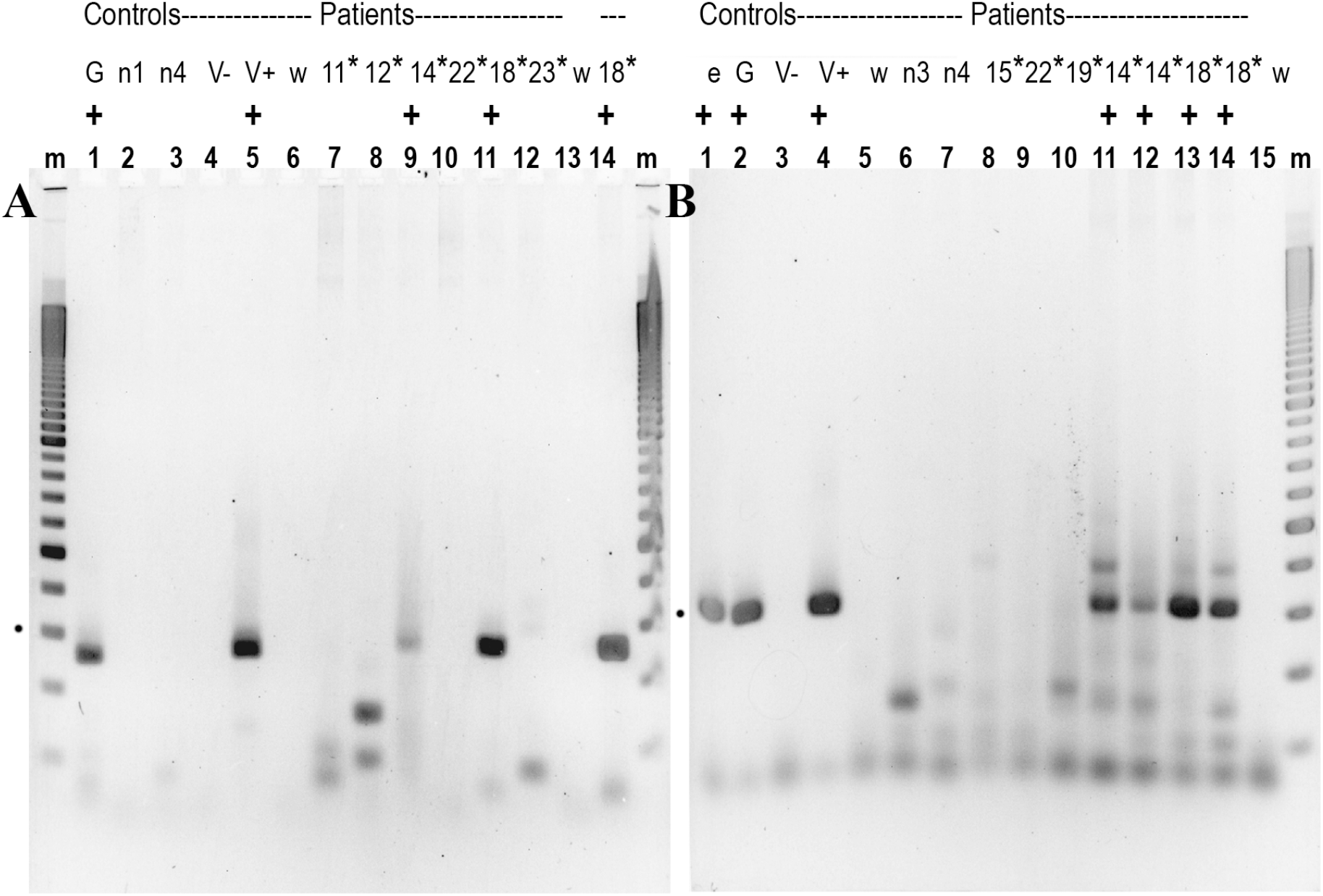
A 2.5% agarose gel showing RT/qPCR products from **A**) the F3/R5 primer synthesized 251bp band and **B**) the 301bp band produced using the F3/R3 primer pair. Positive controls include full length nucleocapsid RNA genome (lanes G) a viral RNA control at e3 copies (lane e), total RNA from CoV19 infected Vero cells (V+ lanes). Negative controls include uninfected Vero cell RNA (V-) and water (w). PBMC uninfected “normal” controls (n1-4 lanes) and representative patient samples numbered with an asterisk (*) are compared. The duplicate 18* lane is from a different aliquot of that patient’s PBMC in another RT/qPCR run (lanes 14-15). Some nonspecific bands are sample specific, e.g., patient 12* (∼150bp band in A) Each duplicate in B is also from a different run with a different aliquot of that extract. A dot is at 300bp in the 100bp ladder marker lanes. SYBR Gold fluorescent DNA is inverted for clarity.

**FIG. 3:**
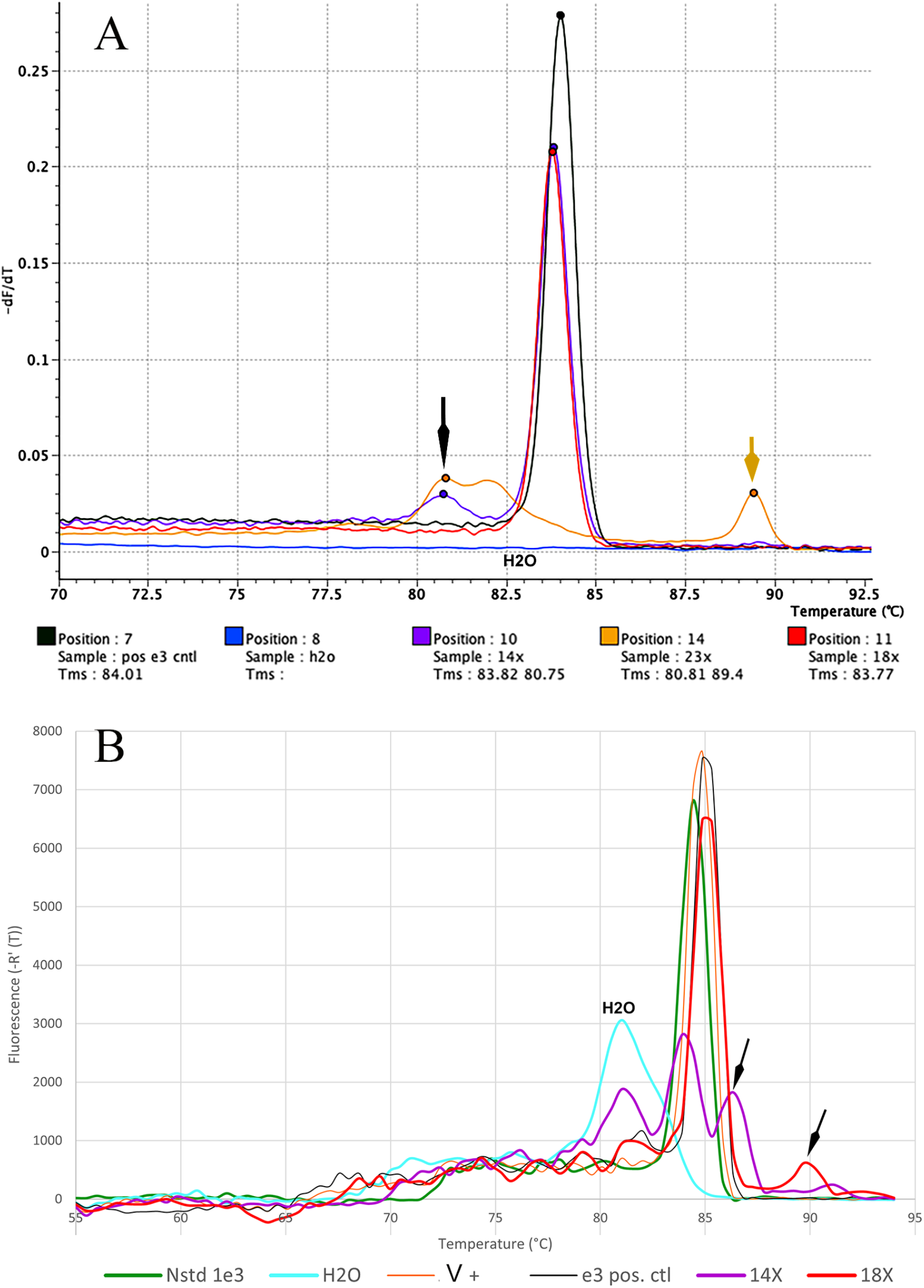
Tm of representative peaks from above gel where **A)** shows the 251bp product. A few smaller peaks with different Tms than the 251bp band could be thresholded. Note the consistency of all NC positive band peaks. The low fluorescent dimer at a Tm at ∼81^°^C is seen in 2 samples (longer black arrow) and a longer faint product, visible on the gel (≥300bp) in 23* is also apparent (short orange arrow) at a Tm of ∼89^°^C. **B**) demonstrates background and non-specific primer dimer peaks, including in the H_2_O control (aqua) that interfered with quantitative Cts. The higher Tm peaks in 14* and 18* on gels looked identical, but had big Tm differences indicating they were sample specific artefact bands.

The two positive patient extracts 14*and 18* that reproducibly yielded the 251bp gel band were also positive for the 301bp band as shown in Fig. 2B. This further supported the presence of longer stretches of CoV19 NC RNA. Each of the positive patient 14* and 18* samples shown in Fig. 2B were also taken from different sample aliquots than those in Fig. 2A. These longer 301bp NC reactions were run in parallel RT/qPCR tubes, with the representative positive and negative controls shown. The CoV19 standards (e, G and V+) all gave relatively clean profiles with only primer-dimer type bands under 100bp. All the control normal PBMC RNAs (n3, n4, and the negative patient samples 15*, 22*and 19*) gave some higher molecular weight smears with a few nonspecific bands under 200bp. Additionally, an ∼ 360bp band is seen in both the stronger 14* sample and the 18* sample (lanes 11 and 14 respectively). This might signify cross-contamination. However, this artefactual peak’s Tm and profile was clearly different in each of these two aliquots.

Fig. 3 shows individual samples with the same nonspecific gel band could be further discriminated from each other by their Tm peaks. The peak profiles of the 251bp (**A**) and 301bp (**B**) RT/qPCR products are representative from the MyGo mini and Stratasys qPCR machines respectively. Duplicates run on both machines also gave comparable results. In Fig. 3A, H_2_O gave a flat negative (blue) line with no peaks whereas all three positives gave the same sharp high fluorescent peak (e+ in black, 14* in purple and 18* in red). The Tm of this peak was the same (83.71±0.6) as previously shown for other positive controls in Fig. 1. Note that although patient 14* showed a weak 251 gel band in Fig. 2A (lane 9), it showed the same dominant Tm peak as the nucleocapsid genome (G), and its Tm peak was clearly much stronger than the nonspecific background. The small artefactual peaks of patient 23* were clearly artefactual by Tm, and were easily thresholded.

Accurate thresholding were more problematic for the longer 301bp product NC primers, (Fig. 3B). In this profile, H_2_O displays an broad nonspecific peak starting at ∼80^°^C and this is clearly due to a primer dimer of ∼50bp. This dimer also is seen in patient samples 14* (purple) and 18* (red) beneath the largest H_2_O peak. The positive 14* sample’s 301bp fluorescent peak lies under the other positive peaks, including the NC control (in green), and has an indistinguishable Tm of 84.19±0.13SD the same as other positive controls in Fig. 1. Another apparently identical nonspecific gel band of ∼350bp is seen in both the 14* and 18* samples (Fig. 2, lanes 11 and 14) yet each of these peaks had clearly different Tms (Fig. 3B, arrows). This confirmed that these 14* and 18* peaks differed in sequence, making cross-contamination of the two samples unlikely. This was borne out by independent sequencing of individual amplified samples of both the 251 and 301bp products in each.

For sequencing confirmation, each of the RT/qPCR bands from positive controls (e, G, V+) as well as both of the bands from the positive patients above (14* and 18*) was determined. All 10 eluted band samples (5 of 251bp and 5 of 301bp) were reamplified, and the bands again gel purified for sequencing. Band eluates run on gels before sequencing showed no other visible bands or primer contamination (data not shown), and all sequences unambiguously confirmed their identity with the NC sequence as detailed in methods. Sequences were confirmed using both strands except DNA close to the F3 and R3 primer ends were read only on one strand. Inspection of run peaks in each confirmed the sequence shown Table I (Methods).

The longer NC sequences amplified here encompass a substantial portion (33%) of the entire nucleocapsid region that codes for a genome-protective protein of 455aa. For future studies, several technical changes can minimize the RT/qPCR artefacts we encountered. First, we had only ∼2e6 total PBMC, optimally yielding only ∼0.6μg RNA for all our repeated RT/qPCR long NC tests. This level was not sufficient for multiple robust triplicate determinations with different primers. Indeed, to ensure reproducible positives we had to use very small samples (1-3ul per reaction with ∼2.5e4 cell RNA/μl). A much higher level of input, e.g., ≥ 5e7 PBMC should reveal more positive patient PBMC. Second, by not rigorously removing genomic DNA, we found more nonspecific bands than in the purer positive RNA controls, such as the viral RNA and cell V+ RNA extracts. DNAse I should be used in the final RNA purification steps. Third, and most importantly, adding a FAM labeled internal amplification reporter probe, rather than nonspecific SYBR Green fluorescence for semiquantitative RT/qPCR assays, should eliminate false positive Cts from primer-dimers and unrelated sequence bands.

## DISCUSSION

The fact that no one else has reported direct reproducible detection of CoV19 RNA sequences in PBMC indicates spread via these cells has not been widely considered even though such cells provide the most fundamental mechanism for disseminating many viruses to distant organs. The use of NC primers extending from the 72bp NC RNA revealed strong and unambiguous evidence that long CoV19 NC sequences were present in 2/11 patient PBMC. The two positive patients (14* and 18*) were validated by sequences that were completely homologous three different positive internal controls, as well as CoV19 NC sequences in the database.

Previous coronavirus experiments using mouse hepatitis virus (MHV), presumed this virus spread by blood because lesions and antigens were distributed in a vascular pattern akin to the one now reproducibly seen in COVID-19 patients. Moreover, after MHV intranasal infection, an infectious viremia with a significant titer was also demonstrated by a colleague here in 1991; the virus was present in whole blood, plasma and buffy coat (white blood cells) 3 days post infection with a titer of 4.1-4.8 logs, LD_50_/ml by mouse bioassay, but by ≥5 days persisted only in buffy coat cells (*14*), and the current study does not rule out very early plasma and PBMC viremia. Early CoV19 blood detection may offer an early opportunity to arrest spread. It is also possible that the relatively low percent of positive patients sampled here at a single early symptomatic time point may underestimate the actual frequency of CoV19 viremia. Experimental MHV viremia was intermittent, and many infectious agents are known to enter the bloodstream in spikes with accompanying intermittent fevers. In any case, the clearance of virus from PBMC remains an important subject, especially for persistent post-infection sequelae. Analyzing PBMC during progressive later stage disease in more susceptible populations such as the elderly, can also be informative for understanding CoV19 viremic clearance and the onset of serious persistent neurological symptoms. Notably, the highest level NC positive patient 18* was 78 years old, and her PBMC was taken 6 days after her symptoms had already developed.

Further studies to determine which PBMC cell type concentrates CoV19 sequences is also critical. While we were limited to a small sample of patient PBMC, with only ∼6e4 cell RNA for 2-3 repeat replicate studies each long NC, it is apparent that only a few select cells (≤1/200) carry these sequences, we presume these are a subset of myeloid cells. Unfortunately, we could not obtain more of our positive patient samples to assess combined in-situ CoV19 antigen/nucleic acid hybridization tests with cell-type specific markers. Nor could we determine if any of the positive cells carry fully infectious CoV19, and not just the nucleocapsid RNA, as by culture or susceptible animal inoculation. Nevertheless, the idea that myeloid lineage circulating cells can be the conduit for dissemination and progressive or chronic disease is linked to their ability to penetrate and take up residence in tissues through the vascular endothelium. Their internalized viral transcripts may also be better protected in brain, and at the same time, as antigen presenting cells, incite inflammatory neurodegenerative changes. Studies of other SARS and MERS coronaviruses have addressed these ACE2 and ACE2-independent monocyte and macrophage pathways (*15*). These coronaviruses also induce a number of ultrastructural membrane modifications in cultured cells that may organize and protect viral assemblies (*16*). The mechanism of CoV19 uptake is likely to be multifactorial in animals with many interrelated cell types, and the huge genetic complexity of coronaviruses should accommodate more than one entry mode. For myeloid cells, uptake of viruses via endocytic membranes with internal sequestration is commonplace, possibly nonspecifically, as a response to recognition of the large CoV19 particle structure, or by myeloid to infected epithelial or endothelial cell attachment. Finally, a selected subset of human migratory myeloid cells in blood have demonstrated Angiotensin converting enzyme 2 (ACE2) that the CoV19 spike protein uses as a receptor for cellular entry (*17*). Therapeutic approaches that target such cells may prevent early organ dissemination of Cov19, in part or whole, and its associated pathological sequelae.

## MATERIALS AND METHODS

### Samples

Collection of de-identified control (“uninfected normal”) blood samples and those from patients with positive CoV19 nasopharyngeal swab tests, first assayed by the US CDC 2019-nCoV_N1 primer–probe set at the Yale-New Haven Hospital (YNHH), were approved by the Institutional Review Board of the Yale Human Research Protection Program (no. FWA00002571, Protocol ID 2000027690). This PBMC project submitted by LM (“Is COVID-19 virus detectable in white blood cells”) was approved by the Yale Biorepository Board of Governors. LM also has HIC approval to study de-identified human normal and pathological blood and tissues (HIC# 8810003391) as well as BSL3 certification.

A total of 18 YNHH peripheral blood monocytes (PBMC) samples were studied. The PBMC were collected and prepared by YNHH nurses, technicians and the Yale IMPACT Research Team from fresh blood and PBMC were separated using standard Ficoll-Paque™ methods, then then frozen at -80^°^C in DMSO-human serum; they had a nominal estimate of 5e6 cells/ml/sample. We selected PBMC samples taken as close to admission as possible to avoid unknown effects of any hospital treatments or progressive disease on viral replication or spread. All patient PBMC samples here were taken within the first week after admission and age range, gender and predisposing risk factors were broadly represented. Patients who subsequently were admitted to the ICU or intubated were excluded.

### Isolation of PBMC RNA

Individual PBMC in DMSO-human serum were thawed briefly in a 37^°^C water bath in the neuropathology biohazard hood BSL2+ lab, mixed gently by pipetting using 1ml pipettor tips, and transferred to 8ml MEM at 22^°^C with only 4-6 samples processed per day. The diluted samples were centrifuged at 700g x 10min at 22^°^C. The NEB Total Miniprep kit was then used as suggested with the following modifications. First, lysis of cells was carefully monitored to minimize breaking large chromatin strands. After discarding each diluted serum-DMSO supernatant, the pellet was tapped to suspend it in residual MEM, 700μl lysis buffer was added and cells allowed to lyse for ∼30” followed by gentle up/down pipetting as needed to disperse lysates using a 1ml tip. Second, only the first “g” column that removes large genomic DNA was used and further DNAse treatment and column purification was avoided because total yields of PBMC RNA could be ≤1μg in 5e6 cells. The g column supernatant then bound to the RNA binding column and eluted in ∼90μl H_2_O; the top ∼65μl away from any residual silica fines or molecular aggregates that were collected and this supernatant recentrifuged. Half (30μl) was used for GAPDH quantification and anti-viral response studies (ML-R and JC-CH) as well as for a standard analysis in triplicate of the CDC 72nt NC primer-reporter assay by the Grubaugh lab (*18*). The other half was divided into subaliquots and frozen for further nucleocapsid (NC) tests. PBMC RNA extracts represented maximally 2.5e4 cells/μl assuming a 100% recovery.

### RNA Standards

A CoV19 nucleocapsid (NC) RNA, a full length CoV19 viral control at e3 from UTMB, and mock infected and CoV19 infected total cell RNA were used as controls. The CoV19 RNA standard for the NC segment was generated as described (*18*) from full-length SARS-CoV-2 RNA (WA1_USA strain of UTMB; GenBank: MN985325) using RT/qPCR followed by T7 transcription from the complementary DNA strand. The FAM dye and BHQ1 quencher were from IDT. This NC RNA, as well as the full length viral RNA, was used for standard dilution comparisons of different real-time qPCR machines used here i.e., BioRad, Stratagene and MyGo mini S (Azura Genomics). Additionally, total cellular RNA from mock-infected and CoV19 infected Vero cells were prepared as described previously by ML-R and JC-CH (*19*). For infection, high-titer stocks of SARS-CoV-2 virus (isolate USA-WA1/2020, CoV-2 WA1) from the BEI reagent repository were obtained by passaging in Vero E6 cells with viral titers determined by plaque assay on Vero E6 cells (ATCC). Virus was cultured exclusively in a biosafety level 3 facility. Cells were infected with virus at a multiplicity of infection (MOI) of 5, and 24 hours post infection were harvested for RNA extraction using the Direct-Trizol RNA kit according to the manufacturer’s instructions (Zymo Research). Mock-infected Vero cell RNA (V-) and CoV19 infected (V+) RNAs were used as paired total cellular RNA controls.

### RT/qPCR

All 18 control and patient PBMC samples were initially tested in triplicate RT/qPCR in the Grubaugh lab (*18*) using the CDC primer/reporter pairs as described and compared to dilutions of the NC RNA and viral e3 RNA as standards. These standards were also used for assays done in duplicate on both Stratasys and MyGo mini qPCR machines. In addition, control mock-infected (V-) and infected (V+) Vero cell RNAs were used as paired standards with longer NC primer pairs. In these, as well as GAPDH RT/qPCR, we used the Luna Universal One-Step RT/qPCR Kit, NEB. The primers GAPDH-F: ACAACTTTGGTATCGTGGAAGG and GAPDH-R: GCCATCACGCCACAGTTTC had a thermal cycle of 55^°^C for 10 minutes for RT, with denaturation at 95^°^C x 1’, then 45 cycles of 95^°^C x 10” and 60^°^C x 30” followed by a standard polishing and melting curve generation at 95^°^C 1’, 55^°^C and 95^°^C x 30”. Relative RNA amounts from patients were normalized to GAPDH of the negative uninfected control PBMC.

The reference sequence for the full NC transcribed segment used for longer NC primer design (LM) was NCBI Reference Sequence: NC_045512.2 from 28274 (ATG) to 29533 with primers F3: TCTTGGTTCACCGC (orange underlined below) and R5: GCAACCATATGCCGTCT yielding a 251bp product. The F3 primer was also used with R3: GATTGCGGGTGCCAATGTG to yield a 301bp sequence reconfirmed the internal sequence of controls and positive patient samples. The relevant portion of the NC is shown with the short 72bp primers and probe in red. The new NC primers and products are shown in other colors starting with the F3 primer (underlined in orange).

Tests of standards with different RT/qPCR temperatures and times gave positive results with the most consistent for the 251bp F3/R5 primer pair at 360nM each was 1) RT at 56^°^C x 15”, 2) 95^°^C x 60” and 3) 45 cycles at 95^°^C x 10”, 58^°^C x10” annealing and 64^°^C x 30” for extension with polishing and melt as for GAPDH above, again using the NEB LUNA One-step RT/PCR kit. The longer 301bp reverse R3 primer with a lower Tm was the same except the 45 cycle phase used 57^°^C x 10” for annealing followed by a 59^°^C x 30” extension. These reactions typically were done in a total of 14μl with 2-3μl PBMC extract to conserve sample for repeat confirmations.

### Purification and sequencing of the NC 251bp and 301bp bands

Each RT/qPCR (0.5 or 1 μl sample) was resolved on mini 2.5% agarose TAE gels, the DNA stained with SYBR gold and the fluorescence imaged digitally in color using a blue emission transilluminator box with an orange barrier filter. Band purification was done as described (*20*). Briefly, the 251bp and 301bp RT/qPCR bands were cut from 2.5% preparative gels (∼5μl sample load), crushed in individual mini weigh boats under parafilm and transferred to 1.5ml polypropylene tubes. Then 200-300μl H_2_O added prior to freeze-thawing and centrifugation at 13,000g for 10’. Eluted supernatants were collected with a 10μl tip to avoid small gel fragments. H_2_O elution/freezing was repeated two more times and SYBR gold stained DNA was followed. More than 70% of the starting band fluorescence appeared in the supernatants while the final residual agarose fluorescence was weak. An aliquot of each eluate was taken for PCR amplification and sequencing using the above NC primers, followed by a second round of band purification. SYBR gold did not interfere with sequencing; some eluates concentrated by lyophilization were washed with 70% EtOH before suspending in 50 μl H_2_O. These had no fluorescence and yielded the identical sequences.

## DATA AVAILABILITY

All data will be findable, fully accessible and freely available under license allowing re-use by any third part for any lawful purpose.

The authors declare no competing financial interests.

## Author contributions

**NP** performed nucleocapsid RT/qRCRs and associated modifications; **ML-R** and **JC-CH** infected Vero Cells, extracted their RNA and did the GAPDH RT/qPCRs of PBMC; **CH** and **NG** did the initial PBMC CDC-72bp assay and quantitation; **LM** designed the project and primers, extracted PBMC RNA, did gel analyses of amplified products and band extraction for sequencing and analysis and wrote the manuscript.

PBMC samples were from the <u>Yale IMPACT Research Team</u>: Kelly Anastasio, Michael H. Askenase, Maria Batsu, Santos Bermejo, Sean Bickerton, Kristina Brower, Molly L. Bucklin, Staci Cahill, Melissa Campbell, Yiyun Cao, Arnau Casanovas-Massana, Edward Courchaine, Rupak Datta, Charles S. Dela Cruz, Giuseppe DeIuliis, Rebecca Earnest, Shelli F. Farhadian, John Fournier, Bertie Geng, Benjamin Goldman-Israelow, Ryan Handoko, William Khoury-Hanold, Christina A. Harden, Akiko Iwasaki, Chaney C. Kalinich, Daniel Kim, Lynda Knaggs, Albert I. Ko, Maxine Kuang, Eriko Kudo, Sarah Lapidus, Joseph Lim, Melissa Linehan, Peiwen Lu, Alice Lu-Culligan, Anjelica Martin, Irene Matos, David McDonald, Maksym Minasyan, Adam J. Moore, M. Catherine Muenker, Maura Nakahata, Nida Naushad, Allison Nelson, Jessica Nouws, Abeer Obaid, Camila Odio, Ji Eun Oh, Saad Omer, Isabel M. Ott, Annsea Park, Hong-Jai Park, Xiaohua Peng, Mary Petrone, Sarah Prophet, Tyler Rice, Kadi-Ann Rose, Lorenzo Sewanan, Lokesh Sharma, Albert C. Shaw, Denise Shepard, Mikhail Smolgovsky, Nicole Sonnert, Yvette Strong, Codruta Todeasa, Maria Tokuyama, Jordan Valdez, Sofia Velazquez, Arvind Venkataraman, Pavithra Vijayakumar, Anne E. Watkins, Elizabeth B. White, Anne L. Wyllie, and Yexin Yang.

## Acknowledgements

We thank Albert Ko for encouraging LM to use Yale repository samples for this study, Arnau Casanovas-Massana who was essential for identifying and gathering early admission PBMC samples, and Shelli Farhadian and Allison Nelson for helping LM with the biorepository application. We are also indebted to the YNHH nurses and technicians who obtained and prepared the PMBC, and also to the many volunteers and patients who agreed to donate to the Yale IMPACT collection. Primer syntheses and sequencing was done by the Yale Keck laboratory. This work was supported by gifts for neurodegenerative research to LM, and the 72bp NC analysis was supported by a Yale School of Public Health startup grant to NDG. We are grateful to Paula Kavathas for her helpful suggestions on the manuscript.

